# Structure and function of the yeast amino acid-sensing SEAC-EGOC supercomplex

**DOI:** 10.1101/2024.10.05.616782

**Authors:** Lucas Tafur, Lenny Bonadei, Yiqiang Zheng, Robbie Loewith

## Abstract

The Seh1-associated complex (SEAC) transduces amino acid signals to the Target of Rapamycin Complex 1 (TORC1), a master regulator of cell growth located on the vacuole membrane. The SEAC acts as a GTPase activating protein (GAP) for Gtr1, a small GTPase that forms a heterodimer with Gtr2, and as part of the EGO complex (EGOC), relays nutrient signals to TORC1. The SEAC is composed of two subcomplexes, SEACIT, an inhibitor of TORC1 that contains the GAP activity, and SEACAT, that has been proposed to regulate the activity of SEACIT. However, molecular details of its regulation are unclear. Here, we determined the cryo-electron microscopy structure of the SEAC-EGOC supercomplex and studied its function in TORC1 amino acid signalling. A single SEAC can interact with two EGOC molecules via SEACIT, binding exclusively to the “active” version of the EGOC as their interaction depends on Gtr1 being loaded with GTP. SEACAT does not modulate nor interact with the EGOC. The GAP activity of the SEAC is essential for the regulation of TORC1 by amino acids, and its loss phenocopies the lack of Gtr1-Gtr2, establishing the SEAC-EGOC complex as an amino acid-sensing hub. Despite being located far from the active site, deletion of Sea2, or its N-terminal β-propeller domain, also results in defects in amino acid signalling to TORC1. We propose that the SEAC-EGOC supercomplex integrates coatomer-like mechanisms of regulation via bidirectional feedback between GAP-GTPase (SEACIT-EGOC) and coat (SEACAT) modules that explain the functional interaction between Sea2 and the GAP activity. Given the conservation between the SEAC and its mammalian ortholog GATOR, we envision that this mechanism is similar in different organisms.

## Introduction

The Target of Rapamycin Complex 1 (TORC1) is a highly conserved master regulator of cell growth that couples nutrient availability with anabolic and catabolic processes. In yeast, amino acids and glucose regulate TORC1 activity through two small Ras-like GTPases, Gtr1 and Gtr2, in a pathway that is conserved to humans. Gtr1 and Gtr2 form a heterodimer that interacts with three other proteins, Ego1, Ego2 and Ego3, together forming the pentameric EGO complex (EGOC)^1^. The EGOC is anchored to the vacuole membrane via the N-terminus of Ego1, which is palmitoylated and myristoylated^2^. Gtr1 and Gtr2 regulate TORC1 activity by changes in their nucleotide loading state: Gtr1^GTP^-Gtr2^GDP^ activates TORC1 under nutrient-replete conditions, while Gtr1^GDP^-Gtr2^GTP^ inactivates TORC1 during starvation^3^. Mechanistically, this involves, at least during changes in glucose levels, the regulated disassembly/assembly of inactive TORC1 helices (TOROIDs)^4^. In mammals, the Rags (RagA/B and RagC/D) are the counterparts of the Gtrs, forming a complex with Ragulator, the counterpart of Ego1-Ego2-Ego3 (Ragulator-Rag)^5^. Like in yeast, RagA/B^GTP^-RagC/D^GDP^ activates and RagA/B^GDP^-RagC/D^GTP^ prevents mTORC1 activation^6,7^, respectively. Changes in the nucleotide loading status of the Rags/Gtrs are coupled to conformational transitions that are thought to regulate the binding to different partners. In mammals, the active Rags can recruit mTORC1 to the lysosome membrane where it can be activated by another GTPase, Rheb^7^.

Gtr1 and Gtr2 have low basal GTPase activity that is stimulated by a dedicated GTPase activating protein (GAP). The Lst4-Lst7 complex acts as GAP for Gtr2^8^, whereas the Seh1-associated complex (SEAC) has specific GAP activity towards Gtr1^9,10^. The SEAC can be functionally subdivided in two subcomplexes: SEACIT and SEACAT. SEACIT, which is composed of Sea1, Npr2 and Npr3, is the subcomplex that harbours the GAP activity of the complex and acts as an inhibitor of TORC1. The activity of SEACIT is believed to be counteracted *in vivo* by SEACAT^9^, which acts as an activator through an unknown molecular mechanism. In mammals, the GATOR complex, which is composed of GATOR1 (SEACIT) and GATOR2 (SEACAT)^11^, and FLCN-FNIP2 (Lst7-Lst4)^12^, act as GAPs for RagA/B and RagC/D, respectively. While specific amino acid sensors that act through GATOR2 have been described^13,14^, as in yeast, how GATOR2 regulates GATOR1 remains a major open question. The currently proposed model of regulation states that when amino acid levels are abundant, proteins that act as amino acid sensors bind their cognate amino acids, which prevents their binding to GATOR2. GATOR2 is then free to inhibit the GAP activity of GATOR1, ultimately activating mTORC1. Conversely, during amino acid starvation, ligand-free amino acid sensors bind GATOR2 and block its inhibitory action on GATOR1, inactivating mTORC1^15^. Although no amino acid sensors have been described in yeast, given their conservation, it is presumed that a similar mechanism of regulation occurs through species-specific sensors^16^.

Both yeast SEACAT and mammalian GATOR2 are composed of five subunits (GATOR2 subunits in parentheses): Sea2 (Wdr24), Sea3 (Wdr59), Sea4 (Mios), Sec13 (Sec13) and Seh1 (Seh1L). Based on secondary structure predictions, initial studies identified a relationship between SEACAT subunits and coatomer complexes^17^ but structural information was lacking. We recently determined the cryo-electron microscopy (cryo-EM) structure of the native yeast SEAC^10^, revealing the interactions between SEACAT and SEACIT. SEACAT forms a central core with similar topology as COPII (SEAC^core^), which serves as an anchor for two flexible wings that are composed of SEACIT (SEAC^wing^). Surprisingly, the active site of SEACIT was not occluded by SEACAT and SEACAT did not inhibit the GAP activity of SEACIT *in vitro*. Contrary to what has been previously suggested^15^, these data are not consistent with SEACAT acting through inhibition of SEACIT. How the SEAC interacts with the EGOC is not known.

Structures of GATOR1^18^ and GATOR2^19^ have been determined individually and show a remarkable conservation with the SEAC, pointing towards a conserved function and regulation. However, compared to the SEAC, where SEACAT and SEACIT form a stable complex, GATOR1 and GATOR2 require an additional complex named KICSTOR to stably interact^20,21^. *In vivo*, GATOR1 interacts with GATOR2 and KICSTOR in an amino acid-insensitive manner^20^. Thus, in cells, both SEACIT/GATOR1 and SEACAT/GATOR2 are constitutively bound and regulate (m)TORC1 as a single complex. Nevertheless, the only structures available of GATOR1 bound to its substrate (Ragulator-Rag) have been determined in the absence of GATOR2 or KICSTOR^18^, precluding a more physiological understanding of the function and regulation of the complex. Furthermore, the role of the GAP activity on TORC1 signalling and its relationship with SEAC function has not been analysed in detail, with only one study in GATOR1 testing a single time point^22^. Therefore, to better understand the molecular mechanism of regulation of the SEAC/GATOR complexes, we set out to determine the structure of the native SEAC bound to the EGOC and characterized its role in amino acid sensing in detail.

## Results

### Cryo-EM structure of the SEAC-EGOC supercomplex

To assemble the SEAC-EGOC supercomplex, we modified slightly our native SEAC purification protocol and incubated the purified SEAC with an excess of recombinant EGOC in the presence of GDP-AlF_3_, which stabilizes GAP-GTPase complexes by mimicking the transition state of the GTP hydrolysis reaction^23^. Despite its relatively small size compared to the SEAC, extra density corresponding to the EGOC was clearly visible in the cryo-EM 2D class averages (Fig. 1a). After 2D and 3D classification steps, we could obtain an overall reconstruction of the complex at 3.2 Å resolution where the core, but not the wings, was well resolved (Extended Data Fig. 1; Extended Data Fig. 2a). To better resolve the wings, we expanded the particles using C2-symmetry, performed local masked 3D classification, subtracted the density of the SEAC^core^ and performed local refinements to obtain a 3.1 Å resolution SEAC^wing^-EGOC reconstruction (Extended Data Fig. 1; Extended Data Fig. 2b). Similarly, to improve the resolution of regions of the SEAC^core^, we performed focused refinements on one monomer and on Sea2-Seh1 and Sea3-Sec13, improving the resolution to ∼3.0 Å (Extended Data Fig. 1; Extended Data Fig. 2c). Finally, to aid in visualization, we combined the resulting maps to create a composite map (Fig. 1b; Extended Data Fig. 2d). We used previous structures of the SEAC^10^ and EGOC^1^ to build a model that includes all 8 SEAC subunits and 5 EGOC subunits, comprising a total of 32 protein chains (Fig. 1c,d; Extended Data Fig. 2e,f).

**Figure 1.**
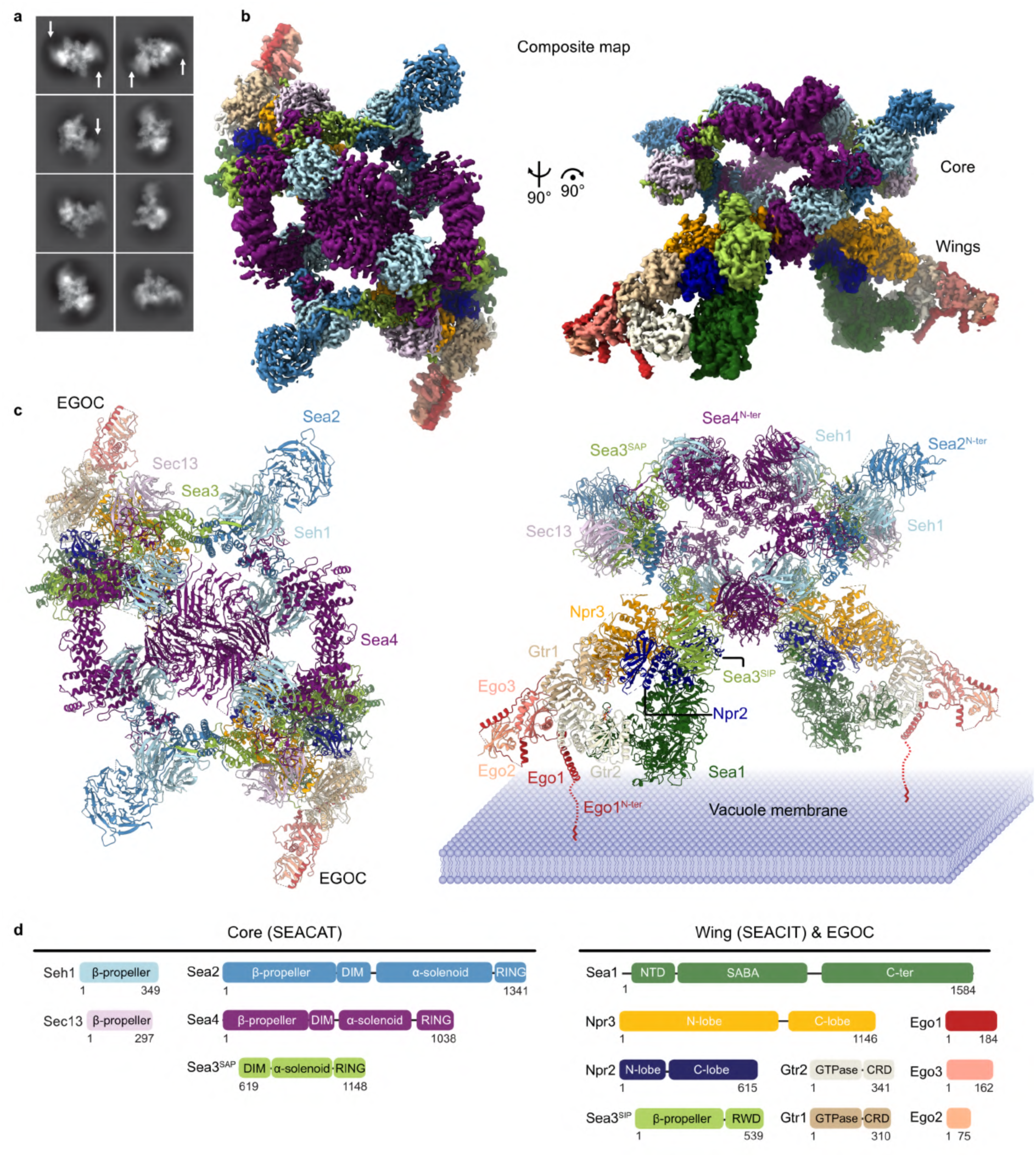
Cryo-EM structure of the SEAC-EGOC super complex. **a**, 2D class averages of final particles used in the reconstruction. Arrows show blurry density corresponding to the EGOC. **b**, Composite cryo-EM map coloured according to subunits. **c**, Model of the SEAC-EGOC super complex. The predicted orientation relative to the vacuole membrane is shown on the right. **d**, Schematic representation of SEAC and EGOC subunits.

The SEAC^core^ is formed by Seh1, Sec13, Sea2, Sea4 and the C-terminal half of Sea3 (SEACAT-portion, Sea3^SAP^), whereas the SEAC^wing^ is formed by Npr2, Npr3, Sea1 and the N-terminal half of Sea3 (SEACIT-portion, Sea3^SIP^). One EGOC molecule is bound to each wing without contacting any element from the core. Accordingly, we observe virtually no change in the overall structure of the SEAC^core^ compared to the unbound SEAC^10^, and only a slight rotation and shift of the SEAC^wings^ (Extended Data Fig. 3a-c). Binding of the EGOC appears to “open” the wings, which rotate around the Npr3-Sea4 interaction (Extended Data Fig. 3c). Superimposition of unbound and EGOC-bound wings reveals that the peripheral and flexible Sea1 is shifted slightly away from Npr2-Npr3 to accommodate binding of the EGOC (Extended Data Fig. 3d). The lack of significant conformational change and the ability of one SEAC to bind multiple EGOC molecules simultaneously is akin to how mTORC1 binds to the Rags^24^ and Rag-TFEB^25^. Curiously, the GAP for Rheb, TSC2, also forms a dimer as part of the Tuberous Sclerosis Complex (TSC)^26^. Thus, the dimeric nature of (m)TORC1 and its regulatory complexes appears to be a recurring feature that likely plays a role in nutrient-sensing by increasing the binding sites to regulatory GTPases. The notable exception appears to be FLCN-FNIP2^27^, which regulates a specific pathway separate from canonical mTORC1 substrates. Nevertheless, like GATOR1^18^, FLCN-FNIP2 appears to have two binding modes^28,29^.

The EGOC is localized to the vacuole membrane^1^, and deletion of *GTR1* and *GTR2* reduces vacuolar localization of the SEAC^10^. Thus, we wished to determine if the complex we observe is compatible with its localization *in vivo*. In our cryo-EM map, Ego1, Ego2 and Ego3 are not resolved as well as Gtr1 and Gtr2 (Extended Data Fig. 2b). However, the last ordered N-terminal helix of Ego1, which is next to the disordered lipidated tail that anchors the complex to the membrane, is sufficiently resolved to determine the orientation of the complex relative to the vacuole membrane (Fig. 1c). In this orientation, the SEAC^core^ faces away from the membrane and is thus accessible for the binding of nutrient sensors, as previously proposed^10^. In turn, the SEAC^wing^ is placed adjacent to the membrane, well-positioning Sea1 for interactions with other proteins like Vps501^30^. This orientation is also consistent with GATOR1 interacting with other lysosomal complexes like KICSTOR^31^ or LYCHOS^32^. Given its flexibility and the lack of interactions between the EGOC and the SEAC^core^, each wing can interact independently with one EGOC molecule. The large degree of freedom of the SEAC^wing^ relative to the SEAC^core^, in addition to the length of the flexible Ego1 N-terminal tail (∼150 Å), could accommodate the binding of the SEAC to one or two EGOC molecules in the context of varying membrane curvatures.

### The EGOC interacts with Npr2 and Sea1

The EGOC interacts extensively with the SEAC^wing^ via Gtr1 and Gtr2 (Fig. 2a). The quality of the Coulombic potential map allowed us to unambiguously assign both nucleotides bound to Gtr1 and Gtr2: whereas Gtr1 is loaded with GDP-AlF_3_ (mimicking GTP), Gtr2 is loaded with GDP (Fig. 2b). Hence, the SEAC-bound EGOC represents the “active” Gtr1^GTP^-Gtr2^GDP^ version.

**Figure 2.**
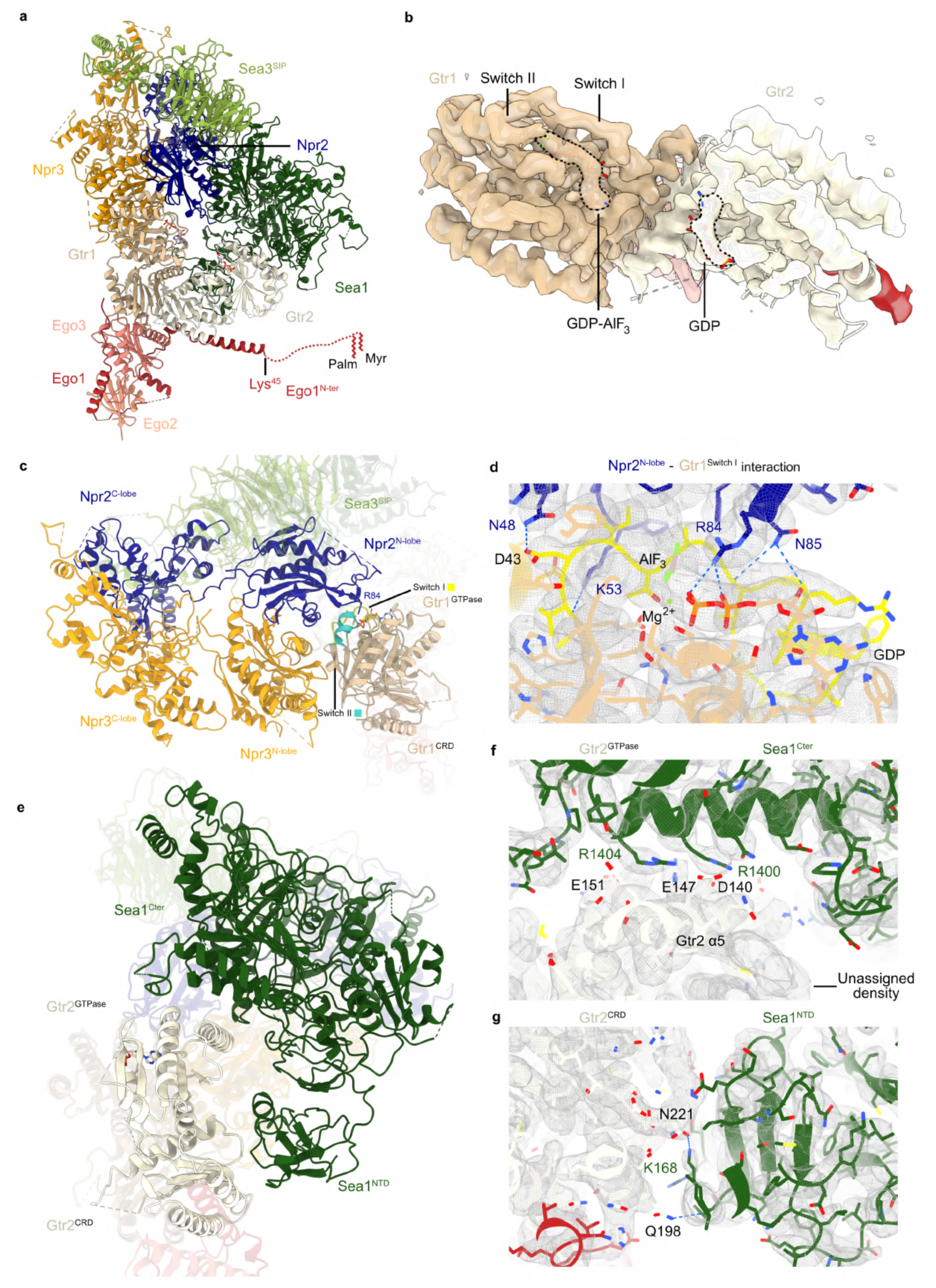
Interactions between the EGOC and SEAC^wing^. **a**, Model of the SEACIT^wing^ bound to the EGOC. The disordered and lipidated tail is indicated schematically. Palm, palmitoylation; Myr, myristoylation. **b**, Cryo-EM density of the Gtr1 and Gtr2 heterodimer. **c**, Binding of Gtr1 to the Npr2^N-lobe^. The Gtr1 Switch I and Switch II regions are indicated. **d**, Interactions between residues near the arginine finger and switch I of Gtr1, coloured yellow. **e**, Binding of Gtr2 to Sea1. **f**, Contacts between the Sea1^C-ter^ and the Gtr1^GTPase^ domains. **g**, Interactions between the Sea1^NTD^ and Gtr2^CRD^ domains.

Two interfaces mediate binding of the EGOC to the SEAC^wing^ and resemble those observed in human GATOR1^18^. The first is formed by Gtr1 and Npr2-Npr3, and the second is formed by Gtr2 and Sea1. Both Npr2 and Npr3 N-lobes (which contain the Longin domain) interact with the Gtr1 GTPase domain, positioning the catalytic “arginine finger” Npr2^R84^ next to GDP-AlF_3_ (Fig. 2c). Insertion of Npr2^R84^ into the nucleotide binding pocket of Gtr1 is stabilized by the presence of the Sea3^SIP^, which sits on top of the Npr2^N-lobe^. Several residues near the catalytic Npr2^R84^ make hydrogen bonds with the switch I region of Gtr1, an element that is only structured when GTP is bound (Fig. 2d). Interestingly, we observe a neighbouring asparagine (Npr2^N85^) next to the arginine finger that interacts with the backbone of switch I, effectively stabilizing its conformation. This asparagine appears to be functionally analogous to the “auxiliary asparagine” observed in RhoGAP^33^ (Extended Data Fig. 4a,b). These interactions ensure that Npr2 only interacts with Gtr1^GTP^.

In contrast to the Npr2-Gtr1 interaction, Sea1 interacts with both the GTPase and C-terminal Roadblock domain (CRD) of Gtr2 on the opposite side of the nucleotide binding pocket (Fig. 2e). In principle, this would mean that binding is not dependent on the nucleotide loading status of Gtr2. The binding interface between Gtr2 and Sea1 is mediated by electrostatic interactions (Extended Data Fig. 4c), which are conserved, to a lesser degree, in the RagC-DEPDC5 (ortholog of Sea1) interaction in GATOR1 (Extended Data Fig. 4d). The Sea1^Cter^ sits on top of the Gtr2 α5 helix (Gtr2^α5^), with two arginine residues (Sea1^R1400^, Sea1^R1404^) close to negatively charged residues (Gtr2^D140^, Gtr2^E147^, Gtr2^E151^) (Fig. 2f), and interactions between the Gtr2^CRD^ and Sea1^NTD^ include two hydrogen bonds (Fig. 2g). Interestingly, we observe an extra density near Gtr2^α5^ that complements the neighbouring β-sheet (Gtr2^β6^) (Fig. 2f). Despite much effort, we were unsuccessful in our attempts to assign this density, but due to proximity, it could correspond to part of the disordered N-terminus of Sea1 (∼100 amino acids are not resolved) (Extended Data Fig. 4e). This extension is not present in the DEPDC5, and, accordingly, no equivalent density is observed in the GATOR1-Rag-Ragulator cryo-EM map (Extended Data Fig. 4f).

Due to the differences in the nature of the interactions in both interfaces, binding appears to be stronger to Gtr1, with Gtr2 serving to sterically restrict the binding to Gtr1 in the context of the Gtr1-Gtr2 heterodimer. This agrees with the lower local resolution of Gtr2 compared to Gtr1 (Extended Data Fig. 2b). Consequently, selectively disrupting the Npr2^N-lobe^-Gtr1^SwitchI^ interface by mutating both the arginine finger and neighbouring asparagine (Npr2^R84AN85A^) is sufficient to remove the SEAC from the vacuole, whereas a single mutation of the arginine finger is not (Extended Data Fig. 4g,h).

In the presence of GDP-AlF_3_, two binding modes have been observed between GATOR1 and Ragulator-Rag: one “GAP-productive” binding as in our structure, and one “non-productive” binding to DEPDC5 (Sea1)^18^. In contrast, we only observe a “GAP-productive” binding mode— we could not obtain a stable complex when incubating the SEAC and EGOC in the presence of the non-hydrolysable GTP analogue GppNHp, which has been used to promote the “non-productive” binding^18^. These results are consistent with the previous proposition that the “non-productive” binding mode is not conserved in yeast^10^. The physiological relevance of the “non-productive” binding remains to be determined.

### The SEAC is unable to bind to the “inactive” EGOC

The Rag and Gtr heterodimers are known to adopt different conformations depending on their nucleotide loading status. We therefore wondered whether, in addition to the “active” Gtr1^GTP^- Gtr2^GDP^ state, other conformations could be accommodated by the SEAC. There is only one previous structure of the EGOC, where both Gtr1 and Gtr2 are bound to GppNHp^1^. Comparison of this structure with ours shows that the Gtrs adopt a very similar conformation despite the different nucleotide loading state, with only a slight shift in Gtr2 due to the binding to Sea1 (Extended Data Fig. 5 a,b). As expected, the conformation of the equivalent states of the Rag heterodimer is similar (Extended Data Fig. 5 c,d). Therefore, both “active” and dually-GTP loaded states would, in principle, be compatible with SEAC binding.

In contrast to the structures described above, there is a drastic conformational change in the RagA-RagC heterodimer in the inactive state (bound to the RagC GAP, FLCN-FNIP2^29^) (Fig. 3a). The shift in RagC^GTP^ compared to RagC^GDP^ widens the distance between GTPase domains from ∼37 Å in the active state to ∼53 Å in the inactive state (Extended Data Fig. 5e). Superimposition of the inactive Ragulator-Rag onto our structure shows that binding of the SEAC to an inactive EGOC is not possible. When bound to GDP, the RagA switch I becomes disordered and thus the interactions with the Npr2^N-lobe^ are lost, and retraction of the switch II produces an additional clash (Fig. 3b). Moreover, the large movement of RagC causes a severe clash with the Sea1^NTD^ (Fig. 3c). A structure of the “inactive” Gtr1^GDP^-Gtr2^GTP^ is lacking, as is a structure of the “inactive” RagA^GDP^-RagC^GTP^ in the absence of an interactor. It is thus possible that the large conformational change observed in the inactive state is induced by FLCN-FNIP2 binding. Therefore, we wished to test our structural predictions. To this end, we took advantage of the fact that deletion of *GTR1* and *GTR2* causes a strong mislocalization of the SEAC to the cytosol^10^ (Fig. 3d,e). By expressing different combinations of Gtr1 and Gtr2 from plasmids, we can test the binding of the SEAC to different versions of the Gtr1-Gtr2 heterodimer in a physiological context by determining if the vacuolar localization of the SEAC is restored. Consistent with our structure, expression of a constitutively “active” EGOC (Gtr1^Q65L^-Gtr2^S23L^) restored normal SEAC localization in *Δgtr1Δgtr2* cells (Fig. 3f), whereas expression of a constitutively “inactive” EGOC (Gtr1^S20L^-Gtr2^Q66L^) did not (Fig. 3g). Thus, the localization differences observed in the different conditions (Fig. 3h) confirm our structural findings and show that the SEAC cannot bind the inactive EGOC *in vivo*.

**Figure 3.**
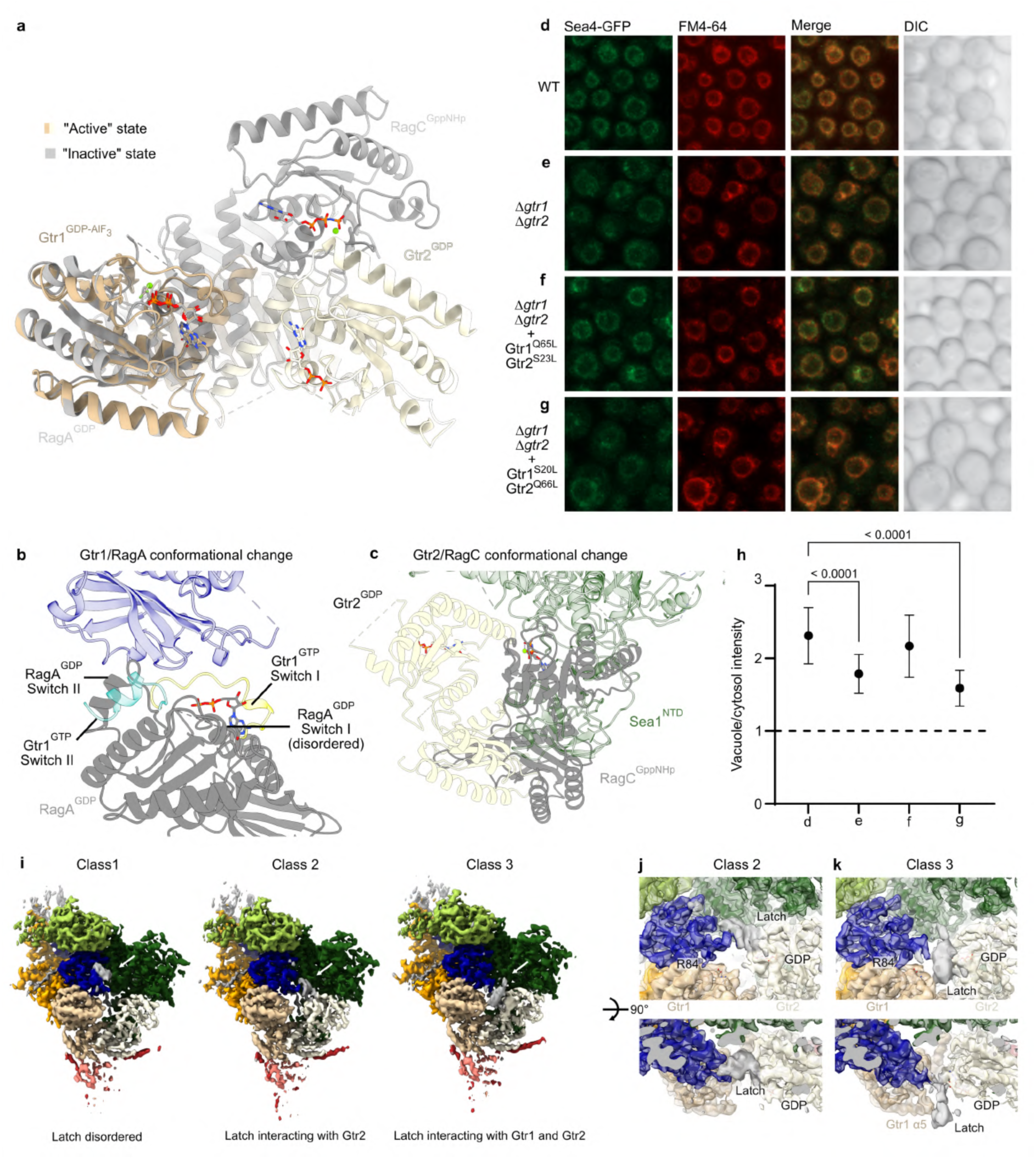
The SEAC binds to the “active”, but not the “inactive”, EGOC. **a**, Comparison between the structure of Gtr1^GDP-AlF3^-Gtr2^GDP^ in the SEAC-EGOC structure (“active”) and the structure of RagA^GDP^-RagC^GppNHp^ when bound to FLCN-FNIP2 (PDB: 6ulg; “inactive”)^29^. **b**, Structural comparison of GTP- and GDP-bound RagA/Gtr1 shows that interactions between Gtr1 and Npr2 are lost in Gtr1^GDP^. **c**, Structural comparison between RagC^GppNHp^ and Gtr2^GDP^ shows that movement of Gtr2 would result in a clash with Sea1. **d**-**g**, Confocal microscopy images showing the localization of the SEAC (using an endogenous GFP tag in Sea4) in the absence (**e**), or with expression of the “active” (**f**) and “inactive” (**g**) versions of Gtr1-Gtr2. **h,** Quantification of the ratio of Sea4-GFP intensity on the vacuole membrane versus the cytosol, in strains shown in panels d-g. The vacuole was labelled using FM4-64. n=60 cells per condition. Mean and standard deviation is shown. Statistical significance is shown with respect to wild type cells. **i**, Cryo-EM maps of the three different classes of the Npr2^latch^. **j**, Zoom-in of the Npr2- Gtr1-Gtr2 interface in Class 2. **k**, Zoom-in of the Npr2-Gtr1-Gtr2 interface in Class 3.

The inability of the SEAC to bind the “inactive” Gtr1-Gtr2 heterodimer is reminiscent of mTORC1, where Raptor can only bind the “active” RagA-RagC heterodimer^34^. Both Raptor and SEAC bind to the Rag/Gtr heterodimer on the same face (Extended Data Fig. 5f,g). In addition, Raptor appears to query the nucleotide loading state of RagC by inserting a stretch referred to as the “claw” between RagA and RagC^34^. The claw would clash with Switch I of RagC if structured and hence, Raptor can only interact with RagC^GDP^. We noted that Npr2 contains an extended loop (∼56 residues) in the vicinity of the arginine finger, which is disordered in our overall SEAC^wing^-EGOC reconstruction. Superimposition of the AlphaFold2 prediction^35^ of Npr2 on our structure revealed that this loop, which we refer to as the “latch”, would be positioned for interactions with the Gtr1-Gtr2 heterodimer in analogous manner as the Raptor claw (Extended Data Fig. 5g). During our analysis of the cryo-EM data by 3D variability^34^, we observed density variation between Gtr1 and Gtr2 GTPase domains that appeared to correspond to part of the latch. Therefore, we performed 3D classification with a mask in this region and successfully resolved, to low resolution, three main classes of the

Npr2^latch^ (Fig. 3i). The first one corresponds to a flexible, disordered state and thus not visible (Class 1). The other two classes show the Npr2^latch^ interacting exclusively with Gtr2 (Fig. 3j; Class 2) or with both Gtr1 and Gtr2 GTPase domains (Fig. 3k; Class 3). In Class3, the Npr2^latch^ inserts into the inter-GTPase domain space and appears to contact the tip of the α5 helix of Gtr1, while shielding the GDP bound to Gtr2. Because the Npr2^latch^ is highly flexible, our densities remain very low resolution and hence we cannot assign specific interactions. The lack of stable and defined interactions, combined with its absence in other species, argues against the Npr2^latch^ being important for SEAC function. Nevertheless, our analysis suggests that different components of the TORC1 signalling pathway in distinct species might reuse similar structural mechanisms to engage with the Gtrs/Rags.

### The GAP activity of the SEAC is required for TORC1 inhibition and reactivation

Based on genetic epistasis results, it has been proposed that SEACAT/GATOR2 regulates (m)TORC1 through modulation of the GAP activity of SEACIT/GATOR1: during nutrient starvation, SEACIT/GATOR1 is active and (m)TORC1 is inhibited, whereas during amino acid repletion, SEACAT/GATOR2 inhibits SEACIT/GATOR1 and activates (m)TORC1^9,11,15^. However, there is no biochemical evidence supporting a direct inhibitory mechanism of SEACAT/GATOR2 on SEACIT/GATOR1. On the contrary, our structure shows that the SEAC^core^ does not block EGOC binding to the SEAC^wing^ nor does it shield the active site, strongly arguing against direct inhibition of catalysis as a mechanism of regulation. Moreover, we previously showed that SEACAT doesn’t inhibit the GAP activity of SEACIT *in vitro*^10^. Therefore, we wished to determine the consequences of abolishing the GAP activity of the SEAC on TORC1 signalling *in vivo*. To this end, we used cells endogenously expressing a catalytically dead SEAC mutant where the arginine finger has been mutated to an alanine (Npr2^R84A^; SEAC^CD^). We have shown that this mutation abolishes the GAP activity but does not affect the integrity of the complex^10^, nor localization to the vacuole (Fig. 2i). As an activity readout, we measured phosphorylation of a TORC1-specific site on the *bona fide* downstream target, Sch9^36^.

Compared to wild type cells, where TORC1 is rapidly (∼2.5 minutes) inactivated and reactivated after removal or addition of amino acids, respectively (Fig. 4a), cells with SEAC^CD^ were almost completely unresponsive to both starvation and repletion during the first 5 minutes (Fig. 4b). We observed a virtually identical response when each individual SEACIT subunit was deleted, or when the interaction between SEACIT and EGOC was disrupted (Npr2^R84AN85A^) (Extended Data Fig. 6). Therefore, abolishing the GAP activity recapitulates the loss of SEACIT, establishing its physiological role in amino acid signalling in yeast. Interestingly, during these time points, amino acid signalling to TORC1 appears to be mediated entirely through the GAP activity of the SEAC, as Δ*gtr1*Δ*gtr2* display the same defects as SEAC^CD^ cells (Fig. 4c). We noted that after this initial phase (< 15 minutes), cells were able to regulate TORC1 to some extent, with activity reaching near wild type levels at around 30 minutes (Fig. 4d,e). This is consistent with the existence of Rag/Gtr-independent pathways^37,38^ that act at later stages of the response (Fig.4f). A two-phase response has been described for TORC1 reactivation by amino acids^39^ and here we show that this also occurs during TORC1 inhibition. Importantly, the SEAC GAP activity appears to be predominant in the first phase, working together with other pathways during the second phase.

**Figure 4.**
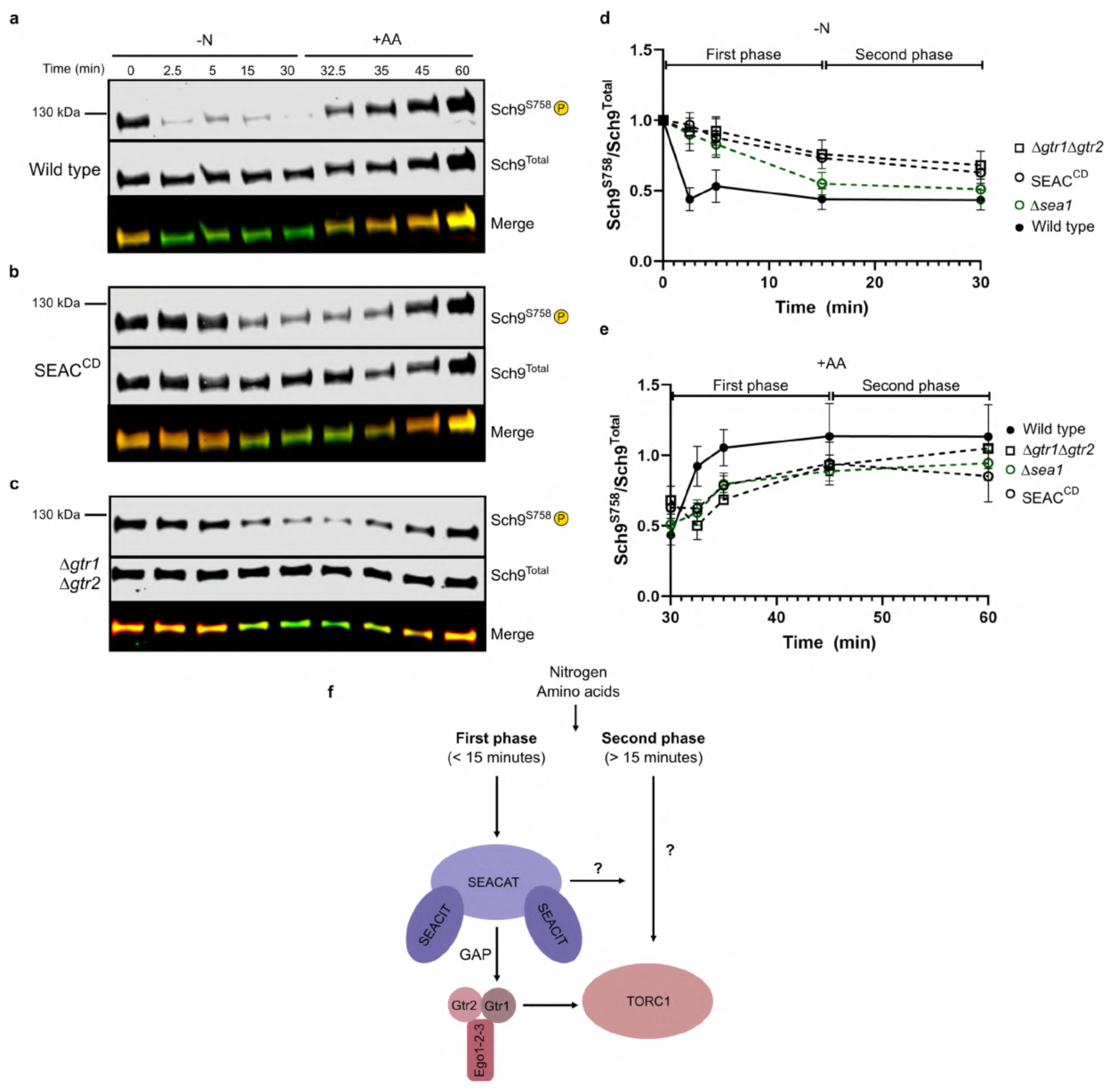
The GAP activity of the SEAC is essential for the regulation of TORC1 by amino acids. **a**, Immunoblot of phosphorylated and total Sch9 after starvation and repletion of amino acids in wild type cells. **b**, Immunoblot of phosphorylated and total Sch9 after starvation and repletion of amino acids in SEAC^CD^ cells. **c**, Immunoblot of phosphorylated and total Sch9 after starvation and repletion of amino acids in Δ*gtr1*Δ*gtr2* cells. **d**, Quantification of relative Sch9 phosphorylation over 30 minutes of starvation in different strains. Data from three independent experiments. **e**, Quantification of relative Sch9 phosphorylation over 30 minutes of repletion in different strains. Data from three independent experiments. λι*sea1* cells are shown for comparison with SEACIT-deficient strains. **f**, Schematic representation of TORC1 regulation by amino acids during the first 30 minutes of starvation and repletion. Two phases are apparent, with the SEAC-EGOC branch predominating on the first (0-15 minutes), while other pathways starting to act during the second (15-30 minutes).

### Sea2 is essential for the function of the SEAC

Previous studies have shown that SEACAT acts as an activator of TORC1^9,10,40^. Consistent with this, we have observed increased rapamycin sensitivity (reflecting lower TORC1 activity) in *Δsea2* and *Δsea4* cells, with a weaker phenotype in *Δsea3* cells^10^. This is likely because Sea3 forms part of both core and wings, as deletion of the Sea3^SAP^ caused a similar phenotype as that of *Δsea2* and *Δsea4* cells^10^. Nevertheless, despite its established role as an activator, no study to date has analysed the individual role of Sea2, Sea3 and Sea4 in acute amino acid signalling; their function has only been inferred from data on GATOR2.

We first asked whether, like its GATOR2 counterparts, subunits forming the SEAC^core^ are necessary for the regulation of TORC1 by amino acids. Given our previous results, we expected a strong reactivation defect in *Δsea2* and *Δsea4* cells, with a much milder phenotype in *Δsea3* cells. Surprisingly, we observed a different phenotype for each strain (Fig. 5a-d). During starvation, *Δsea4* (Fig. 5b) and *Δsea3* cells (Fig. 5c) behaved almost like wild type cells, having an impaired response only at the first time point and quickly recovering afterwards. Interestingly, *Δsea2* cells displayed SEACIT-like kinetics, with impaired first phase response and slower recovery during the second phase (Fig. 5d,e). Greater differences between strains were apparent during amino acid repletion, with *Δsea2* cells being completely unresponsive to amino acids in the first phase, *Δsea3* cells being unresponsive only at the first time point, and *Δsea4* cells responding normally (Fig. 5f). Furthermore, *Δsea2* cells had very low TORC1 activity (Fig. 5d) and a slow growth rate (data not shown). Thus, contrary to previous suggestions^40^, SEACAT subunits appear not to be redundant. Comparison of the different mutants revealed that the TORC1 activity kinetics of *Δsea2* cells overlapped better with SEAC^CD^ cells than with *Δsea3* and *Δsea4* cells (Fig. 5e,f), suggestive of a Sea2-GAP axis for amino acid signalling. Therefore, as in GATOR2^19^, Sea2 plays an essential role in the SEAC. Contrary to our results, loss of Mios (Sea4) in GATOR2 causes defects in amino acid signalling^19,41^. However, deletion of Mios results in degradation of Wdr24 (Sea2)^19,21,41^, making the interpretation of these results difficult.

**Figure 5.**
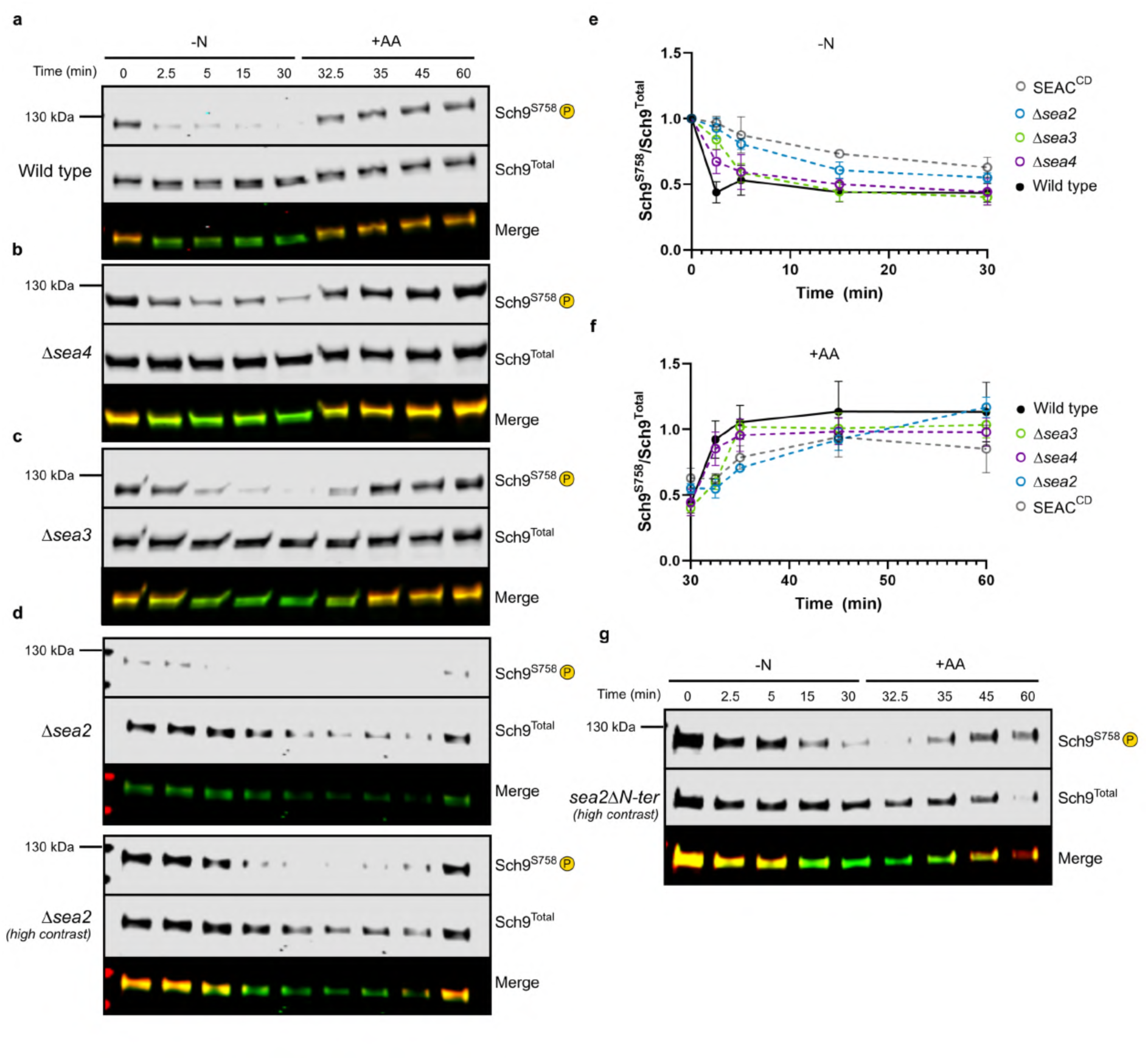
Sea2 is essential for the function of the SEAC. **a**, Immunoblot of phosphorylated and total Sch9 after starvation and repletion of amino acids in wild type cells. **b**, Immunoblots of phosphorylated and total Sch9 after starvation and repletion of amino acids in Δ*sea4* cells. **c**, Immunoblots of phosphorylated and total Sch9 after starvation and repletion of amino acids in Δ*sea3* cells. **d**, Immunoblots of phosphorylated and total Sch9 after starvation and repletion of amino acids in Δ*sea2* cells. These cells have a lower overall TORC1 activity and slow growth. **e**, Quantification of relative Sch9 phosphorylation over 30 minutes of starvation in wild type and SEACAT deletion strains. Data from three independent experiments. **f**, Quantification of relative Sch9 phosphorylation over 30 minutes of repletion in wild type and SEACAT deletion strains. Data from three independent experiments. **g**, Immunoblot of phosphorylated and total Sch9 after starvation and repletion of amino acids in the *sea2*Δ*N-ter* strain. Quantification overlaps with Δ*sea2* cells and is omitted from panels e and f for clarity.

In both the SEAC^core^ and GATOR2^19^, the N-terminal β-propeller of Sea2 (Sea2^N-ter^, Wdr24^N-ter^ in GATOR2) protrudes outwards of the complex, is highly flexible (Extended Data Fig. 2a), and is the only β-propeller that does not interact with any other subunit (Fig. 1). Recent studies have found that the Wdr24^N-ter^ is subject to nutrient-responsive post-translational modifications^42,43^. Moreover, in addition to serving as a binding platform for the leucine sensor Sestrin2^19^, it is also essential for GATOR2 function^19^. Thus, we reasoned that the Sea2^N-ter^ plays a similar role in the SEAC. In agreement, deletion of the Sea2^N-ter^ caused the same defects observed in Δ*sea2* cells (Fig. 5g), showing that, like in GATOR2, this region mediates an essential function of the complex.

Compared to Sea3 and Sea4, Sea2 does not interact with the SEAC^wing^ and is located far from the active site, with the Sea2^N-ter^ being at least ∼180 Å away from Npr2^R84^. Therefore, it is unlikely that Sea2 acts directly on Npr2. Rather, data are consistent with the GAP activity and Sea2 acting in parallel to regulate TORC1. To probe this idea, we generated *sea2Δ npr2^R84A^* (SEAC^CD^Δ*sea2*) double mutant cells in which we assessed TORC1 activity following amino acid starvation and subsequent repletion. As in the single mutants, the first phase response was blunted during both starvation and repletion (Fig. 6a-c). Intriguingly, SEAC^CD^Δ*sea2* cells displayed a normal second phase response, with TORC1 activity reaching wild type levels at 15 minutes (Fig. 6a-c). To further test the relationship between the GAP activity and Sea2, we performed growth assays on rapamycin. Here, Δ*sea2* cells displayed severe rapamycin sensitivity, as previously observed^10^, whereas SEAC^CD^ cells were rapamycin resistant (Extended Data Fig. 6f). This matches the expected overall lower TORC1 activity with the deletion of *SEA2* and higher TORC1 activity when abolishing the GAP activity. Contrary to the effects on acute amino acid signalling, where combining the two mutations resulted in a different phenotype than individual mutants, SEAC^CD^Δ*sea2* cells displayed similar resistance to rapamycin as SEAC^CD^ cells (Extended Data Fig. 6f). Therefore, the GAP activity appears to be epistatic to Sea2 for the regulation of overall TORC1 activity, whereas there is a more complex functional relationship between the two interfaces during acute amino acid signalling. Combined with our observations in SEACAT deletion strains, our results show that the effects on acute regulation of TORC1 by amino acids are not reflected by growth assays on rapamycin.

**Figure 6.**
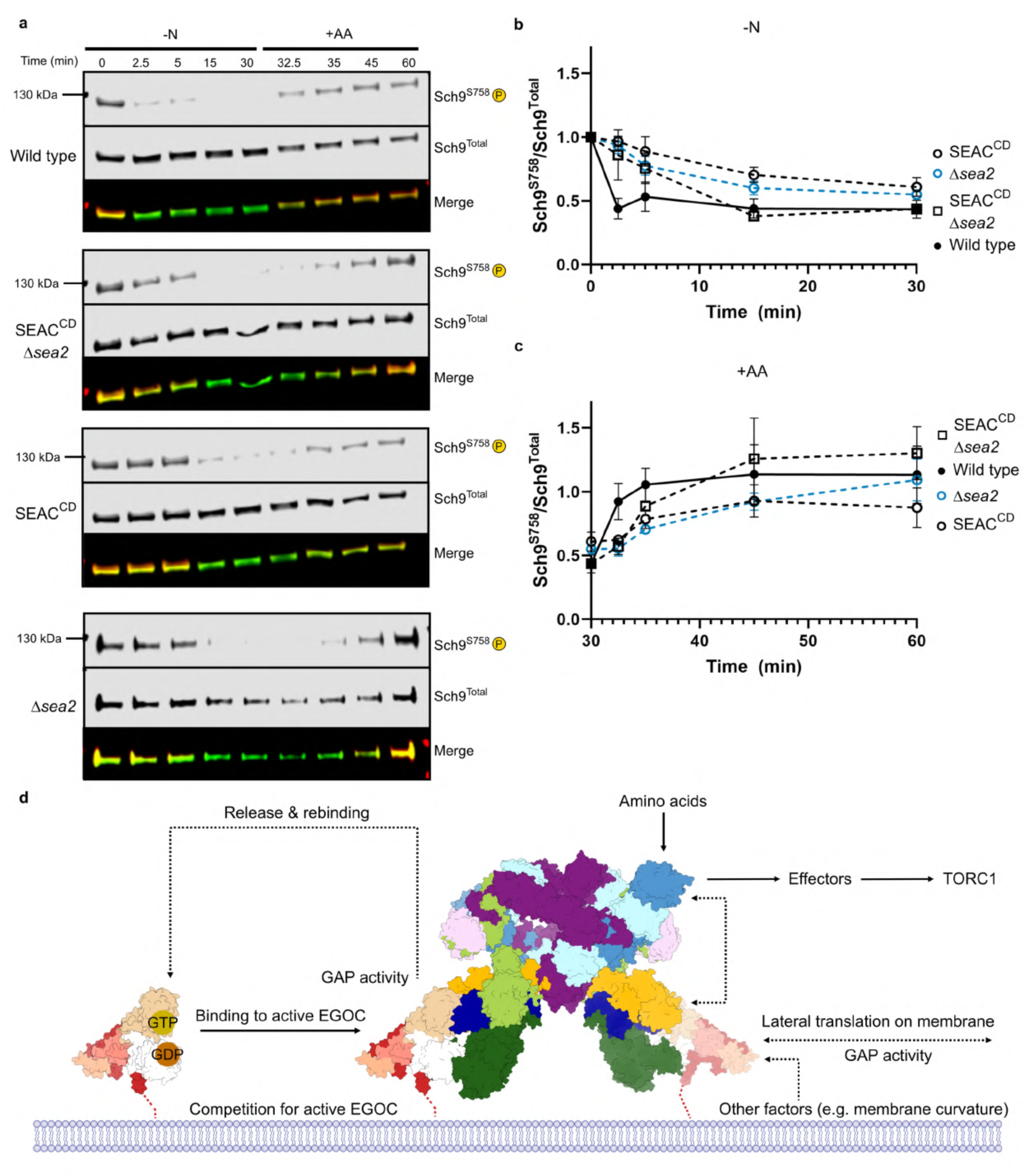
Bidirectional signalling within the SEAC. **a**, Immunoblots of phosphorylated and total Sch9 after starvation and repletion of amino acids in wild type, SEAC^CD^Δ*sea2*, SEAC^CD^ and Δ*sea2* cells. **b**, Quantification of relative Sch9 phosphorylation over 30 minutes of starvation in wild type, SEAC^CD^Δ*sea2*, SEAC^CD^ and Δ*sea2* cells. **c**, Quantification of relative Sch9 phosphorylation over 30 minutes of repletion in wild type, SEAC^CD^Δ*sea2*, SEAC^CD^ and Δ*sea2* cells. Data from three (wild type and SEAC^CD^) and two (SEAC^CD^Δ*sea2*) independent experiments. **d**, Model of regulation of the SEAC on the vacuolar membrane. We propose that the GAP activity regulates the localization of the complex and Sea2, which transmits signals to TORC1 via protein effectors. Selectivity for the active EGOC would also result in competition with TORC1 and the control of basal TORC1 activity. Other factors that affect membrane properties may also regulate the core by affecting the GAP activity or Gtr1 GTP hydrolysis.

### Model of regulation of SEAC function

We propose a model by which the GAP activity, by producing Gtr1^GDP^, regulates the distribution and dynamics of the SEAC along the vacuole membrane (Fig. 6d). As the SEAC cannot interact with Gtr1^GDP^ (Fig. 2h; Fig. 3g), each wing would quickly dissociate from the EGOC after catalysis. This is reminiscent of COPII, where the GAP activity is coupled to dissociation of the coat^44^. In the SEAC, given that both GAP and coat modules are part of the same complex, this would result in the dynamic association of the SEAC with EGOC molecules on the vacuole membrane without dissociation of the GAP-coat complex. Using the wings, the SEAC would then move along the vacuole through Gtr1 GTP hydrolysis. The preference of the SEAC for the active state of the Gtr1-Gtr2 heterodimer may potentially result in competition with TORC1 for the active EGOC, additionally regulating basal TORC1 activity. Importantly, mobilization of the SEAC by the GAP activity would also regulate the localization of the Sea2^N-ter^, functionally linking these two interfaces. This could explain why localizing the SEAC to the vacuolar membrane is not sufficient for TORC1 regulation (as SEAC^CD^ cells display grossly normal SEAC localization); normal signalling likely requires the redistribution of Sea2 to specific vacuolar membrane domains. Of note, the vacuolar membrane is known to phase separate into distinct lipid domains after long periods of starvation^45^.

Our results extend the structural relationship of the SEAC with coatomer complexes and suggest that molecular mechanisms observed in other systems, such as COPI and COPII, might equally function in the SEAC. In this manner, the architecture of the SEAC-EGOC supercomplex is suited for bidirectional regulation between coat (SEAC^core^) and GAP-GTPase (SEAC^wing^-EGOC) modules. For example, in COPI, membrane curvature promotes coat disassembly through modulation of the activity of Arf1GAP on the Arf1 GTPase^46^. Something similar might occur on the vacuole, where changes in membrane curvature or lipid packing could trigger GTP hydrolysis on Gtr1 and SEAC dissociation. Other factors that interact with EGOC/Ragulator might then transmit nutrient cues to the SEAC^core^ by physically affecting the membrane or the nucleotide loading status of the Gtrs/Rags. Such a mechanism could couple signals from inside and outside the lysosome/vacuole, explaining, for instance, the role of the vacuolar ATPase in amino acid sensing^47^. Therefore, the GAP activity is at the centre of the regulation of the SEAC because it transduces signals from and to the core, and, in particular, to Sea2. How does the Sea2^N-ter^/Wdr24^N-ter^ regulate (m)TORC1 is an important question for future study. It is likely that the Sea2^N-ter^/Wdr24^N-ter^ interacts with additional protein effectors to exert its regulatory role; sensors like Sestrin2 and modifications that have been reported to act on this region (phosphorylation^42^ and methylation^43^) might therefore impair or promote these interactions.

In conclusion, we propose a bidirectional model of regulation between SEAC and EGOC that integrates amino acid signalling to TORC1.

## Methods

### Yeast strains

Yeast strains used in this study are of TB50 background. Classical recombination-based or CRISPR-based techniques were used to construct different strains. All strains were verified by PCR, sequencing or Western Blot (where applicable). Yeast strains used in this study are listed in Extended Data Table 1 and plasmids in Extended Data Table 2.

### Native SEAC purification

Native SEAC was purified using a TAP tag in Sea4, essentially as previously described^10^, with the only modification being that the complex was eluted in EGOC binding buffer mixed with TEV protease (20mM HEPES pH 7.4, 150mM NaCl, 5mM DTT, 0.8g/l EGOC, 1mM GDP, 2mM MgCl_2_, 2mM AlCl_3_, 20mM NaF). The complex was prepared freshly for cryo-EM grid preparation.

### EGOC purification

Recombinant full-length EGOC was purified as previously described^10^. Plasmids encoding wild type Gtr1 and Gtr2 were cotransformed with a plasmid containing full length Ego1, Ego2 and Ego3, where Ego1 was N-terminally tagged with a histidine tag.

### Cryo-EM grid preparation

5μl of SEAC-EGOC mix was deposited on 1.2/1.3 Quantifoil Au grids and plunge frozen in a Leica GP2 instrument.

### Cryo-EM data collection

Grids were screened for quality, and preliminary data collection was performed on a 200 kV Talos Arctica equipped with a Falcon 3 direct electron detector (DCI-Geneva).

Final data were collected on a 300 kV Titan Krios equipped with a Selectris X energy filter and a Falcon 4i direct electron detector in the Dubouchet Center for Imaging (DCI-Lausanne). 11,177 untilted and 7,253 35° tilted movie frames were collected with a total exposure of 40e^-^/ Å^2^ and a nominal pixel size of 0.726 Å.

### Cryo-EM data processing

The cryo-EM processing pipeline is detailed in Extended Data Fig. 1b. Movies in EER format were pre-processed (Patch Motion correction and Patch CTF estimation) on-the-fly with CryoSPARC Live. All subsequent steps were performed in cryoSPARC^48^. Ultimately, from 1,649,067 template picks, we selected 208,039 well-aligning SEAC-EGOC particles, that were refined with C2-symemtry to 3.2 Å resolution (consensus map). To improve different regions of the complex, particles were symmetry-expanded, particle subtracted (to eliminate signal from the other monomer and wings) and subject to local (with non-uniform regularisation) refinements on different regions of the core. After local refinement on the wing (which was not resolved in the consensus map), focused 3D classification on this region yielded two classes with good EGOC density. We performed an additional 3D classification with high resolution threshold to select better-aligning particles, signal subtracted the rest of the SEAC and obtained a reconstruction at 3.1 Å resolution from 133,077 particles. We noted weak density between the Gtr1 and Gtr2 GTPase domains that could correspond to a disordered loop of Npr2 (Npr2^lid^; Extended Data Fig. 5d). We therefore used a mask to perform focused 3D classification on this region and obtained three different reconstructions with variations in this density, which we called States 1 to 3. We individually locally refined each state.

All maps (except the three Npr2^lid^ states) were sharpened using DeepEMhancer^49^. To create a composite map, all focused maps were aligned in UCSF Chimera^50^ to the consensus map, regions around 4 Å were selected and merged using the “vop maximum” command. Local resolution was estimated with Blocres^51^ within cryoSPARC. Figures were made using UCSF ChimeraX^52^.

### Model building

The structure of the SEAC (PDB:8ADL) was used as a starting model. For the EGOC, AlphaFold2^53^ predictions and a published crystal structure (PDB:6JWP)^1^ were used as starting models. Initial rigid-body fitting was performed in Chimera and subsequent manual building was performed in Coot^54^. Models were real space refined iteratively in Phenix^55^. Some visible stretches of density were not built due to our inability to unambiguously assign those densities. In addition to those around Sea1 and Gtr2 (Extended Data Fig. 4e), there is another density in the cavity between Npr2-Npr3-Sea3^SIP^. Attempts to use automated model building tools were also unsuccessful. Thus, we preferred to not interpret them.

### Immunoblots

Amino acid starvation and repletion experiments were performed as follows. Yeast was grown to an OD_600_ of 0.8-1 in complete synthetic media (CSM; 2% glucose, yeast nitrogen base without amino acids with ammonium sulfate, and drop-out mix). After taking the first time point (0), cultures were filtered on 0.2 μm filter membranes and resuspended in an equal volume of CSM -amino acids -ammonium sulfate (2% glucose, yeast nitrogen base without amino acids nor ammonium sulfate). After 30 minutes, drop-out mix was added to the culture at 1X final concentration.

For each time point, 10 mL of culture was mixed with cold TCA (final concentration 6%) and kept on ice. Cells were pelleted, washed with acetone and dried on a SpeedVac. Dried samples were processed or kept at −80°C. Cells were lysed using a bead beater in lysis buffer (25 mM Tris pH 6.8, 6M Urea, 1% SDS) and mixed with 2X SDS-sample buffer (25 mM Tris pH 6.8, 20% glycerol, 2% SDS, 0.02% bromophenol blue, 200 mM DTT). Samples were resolved on a 4-20% gradient TGX gel (Bio-Rad) and transferred to nitrocellulose membranes on an iBlot 2 (ThermoFisher). Membranes were blocked with 1X TBS buffer with 5% BSA for 30-60 minutes and incubated with primary antibodies overnight at 4°C (1:5000, anti-Sch9; 1:25000 anti-Sch9^S758^). After washing with 1X TBS, secondary antibodies were incubated for 1 hour at room temperature and the blot was imaged on a LiCOR system. Quantification was performed on ImageStudio 5.5 (LiCOR) and plotted and analysed using GraphPad Prism.

### Confocal microscopy

Strains were grown on CSM to an OD_600_ of about 0.7–1. Z-stacks were collected on a Zeiss LSM800 confocal laser scanning microscope and processed using ImageJ 1.52p. The ratio of vacuole-to-cytosol GFP intensity was essentially done as previously^10^, using FM4-64 to delineate the vacuole membrane. Statistical analysis was performed in GraphPad Prism using Brown-Forsythe and Welch ANOVA tests.

## Data availability

Cryo-EM maps and models have been deposited in the Electron Microscopy Database (EMDB) and Protein Data Bank (PDB), respectively, with accession codes: EMD XXX & PDB XXX for the full SEAC-EGOC, and EMD XXX & PDB XXX for the SEAC^wing^-EGOC.

---

**Table 1.** Map and model refinement statistics.

|  | SEAC-EGOC | SEAC<sup>wing</sup>-EGOC |
| --- | --- | --- |
|  | (EMD-XXX) | (EMD-XXX) |
|  | (PDB XXX) | (PDB XXX) |
| **Data collection and processing** |  |  |
| Magnification | 165,000 |  |
| Voltage (kV) | 300 |  |
| Electron exposure (e<sup>-</sup>/Å<sup>2</sup>) | 40 |  |
| Defocus range (μm) | -1.6 to -0.6 |  |
| Pixel size (Å) | 0.726 |  |
| Symmetry imposed | C2 |  |
| Initial particle images (no.) | 1,649,067 |  |
| Final particle images (no.) | 208,039 | 133,077<br>(symmetry-expanded) |
| Map resolution (Å) | 3.2 | 3.1 |
| FSC threshold | 0.143 | 0.143 |
| Map resolution range (Å) | 2.207-41.635 | 2.207-43.895 |
| **Refinement** |  |  |
| Initial model used (PDB code) | 8adl, 6jwp | 8ae6 |
| Model resolution (Å) | 4.0 | 3.4 |
| FSC threshold | 0.5 | 0.5 |
| Model composition |  |  |
| Non-hydrogen atoms | 114312 | 27412 |
| Protein residues | 14138 | 3355 |
| Nucleotides | 4 | 2 |
| Ligands | ZN: 28 | MG: 1 |
|  | MG: 2 | AF3: 1 |
|  | AF3: 2 |  |
| B factors (Å<sup>2</sup>) | (min/max/mean) | (min/max/mean) |
| Protein | 0/688.50/118.99 | 17.42/197.63/80.27 |
| Nucleotide | 58.26/112.79/90.99 | 26.49/44.56/35.52 |
| Ligand | 71.78/192.70/134.95 | 61.03/74.73/68.08 |
| R.m.s. deviations |  |  |
| Bond lengths (Å) | 0.004 | 0.003 |
| Bond angles (°) | 0.977 | 0.553 |
| Validation |  |  |
| MolProbity score | 1.49 | 1.53 |
| Clashscore | 4.27 | 4.46 |
| Poor rotamers (%) | 0.00 | 0.00 |
| EMRinger score | 1.89 | 2.45 |
| Ramachandran plot |  |  |
| Favored (%) | 95.94 | 95.59 |
| Allowed (%) | 4.06 | 4.41 |
| Disallowed (%) | 0 | 0 |
| Ramachandran Z-score |  |  |
| Whole | -0.98 | -0.98 |
| Helix | -0.19 | -0.15 |
| Sheet | -0.19 | -0.42 |
| Loop | -1.13 | -1.08 |

**Extended Data Figure 1.**
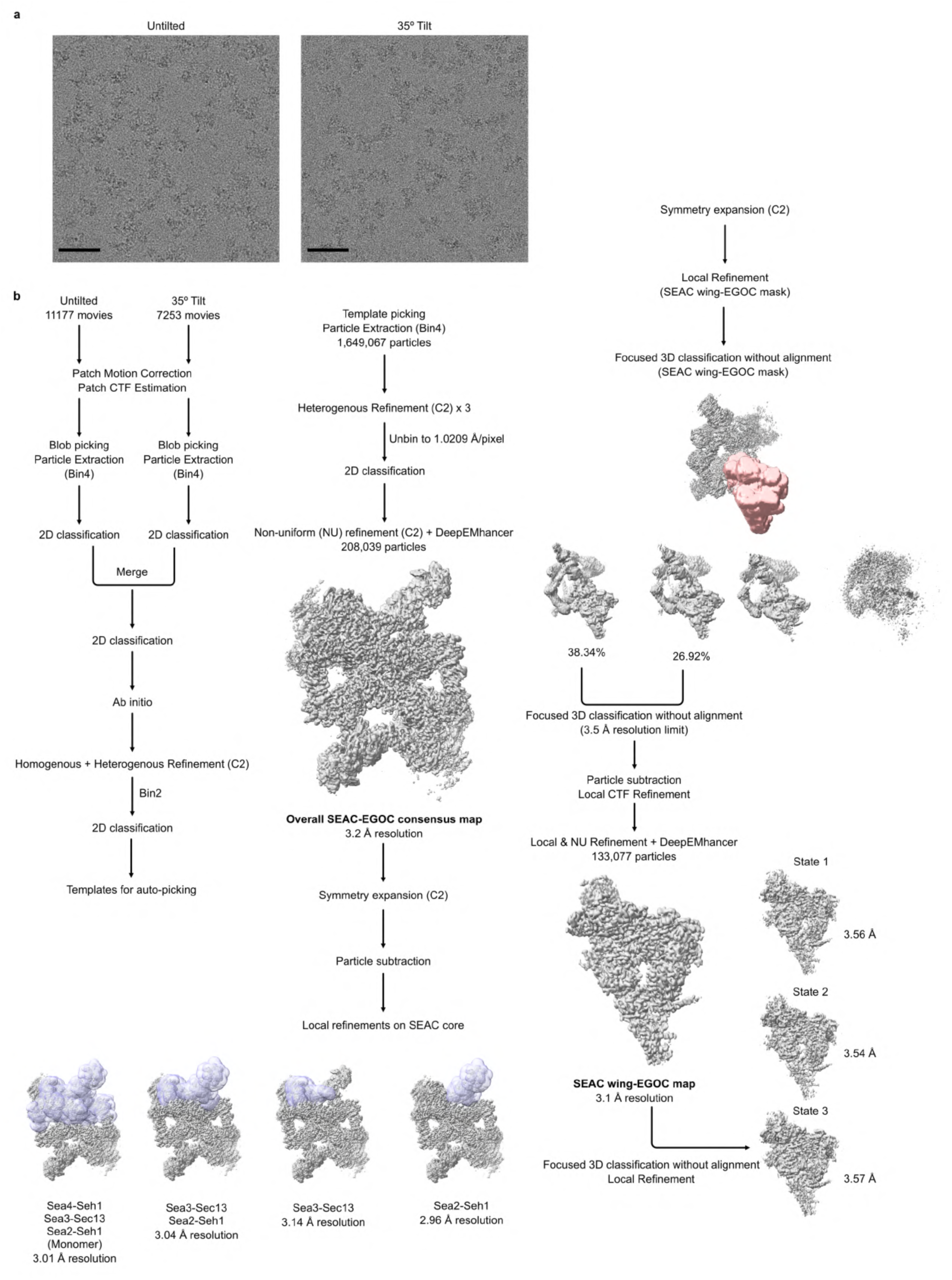
Cryo-EM data processing pipeline. **a**, Representative micrographs from untilted and tilted datasets. Scale bar = 50 nm **b**, Processing pipeline for the analysis of cryo-EM data.

**Extended Data Figure 2.**
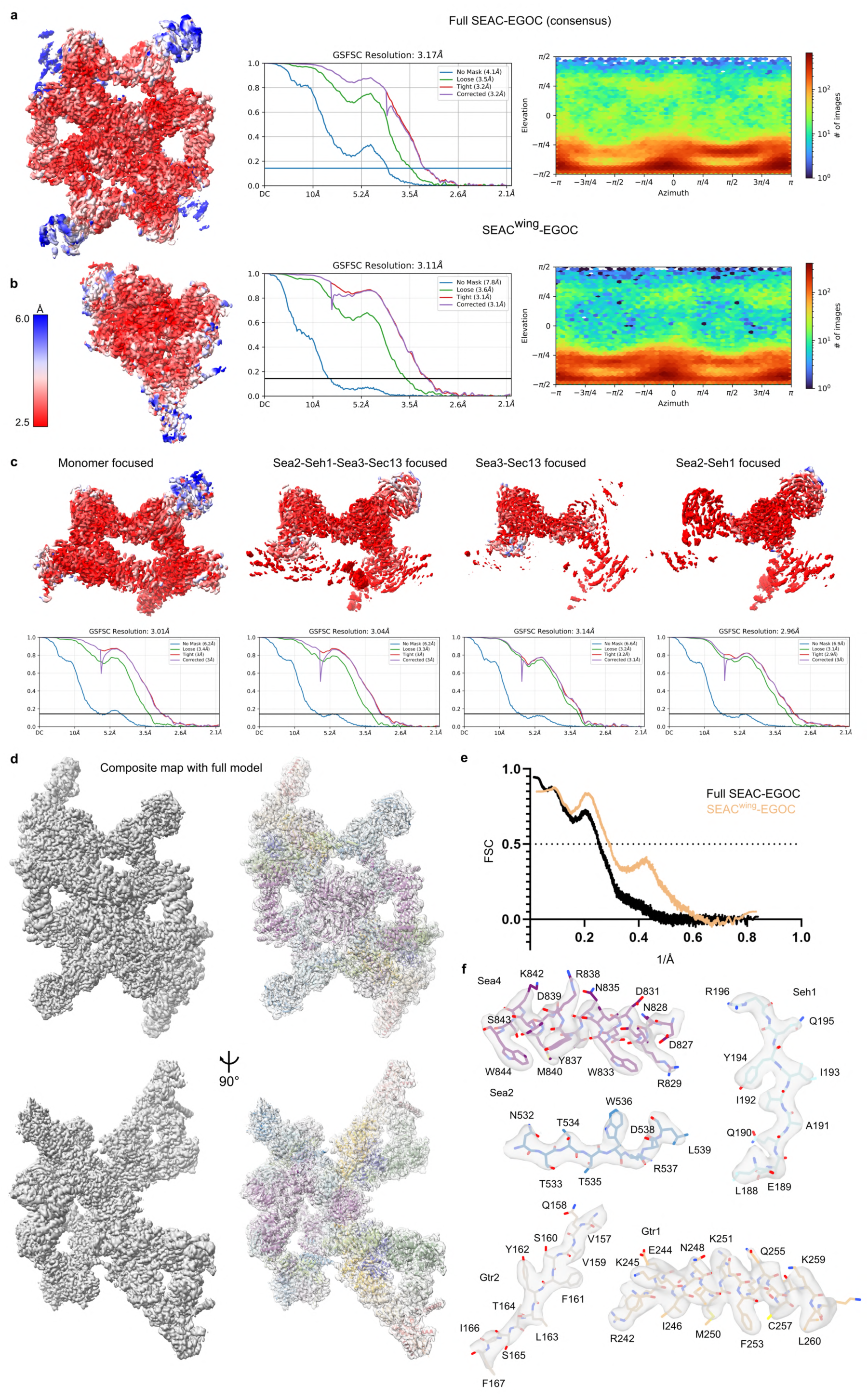
Map and model validation statistics. **a**, Local resolution estimate, half-map FSC plot and orientation distribution from the final refinement of the full consensus reconstruction. **b**, Local resolution estimate, half-map FSC plot and orientation distribution from the final refinement of the SEAC^wing^-EGOC reconstruction. **c**, Local resolution estimates and half-map FSC plots for the focused refinements on the SEAC^core^. **d**, Composite map fitted with full SEAC-EGOC model. **e**, Model-map FSC plots for the Full SEAC-EGOC and the SEAC^wing^-EGOC. **f**, Representative densities for the cryo-EM maps.

**Extended Data Figure 3.**
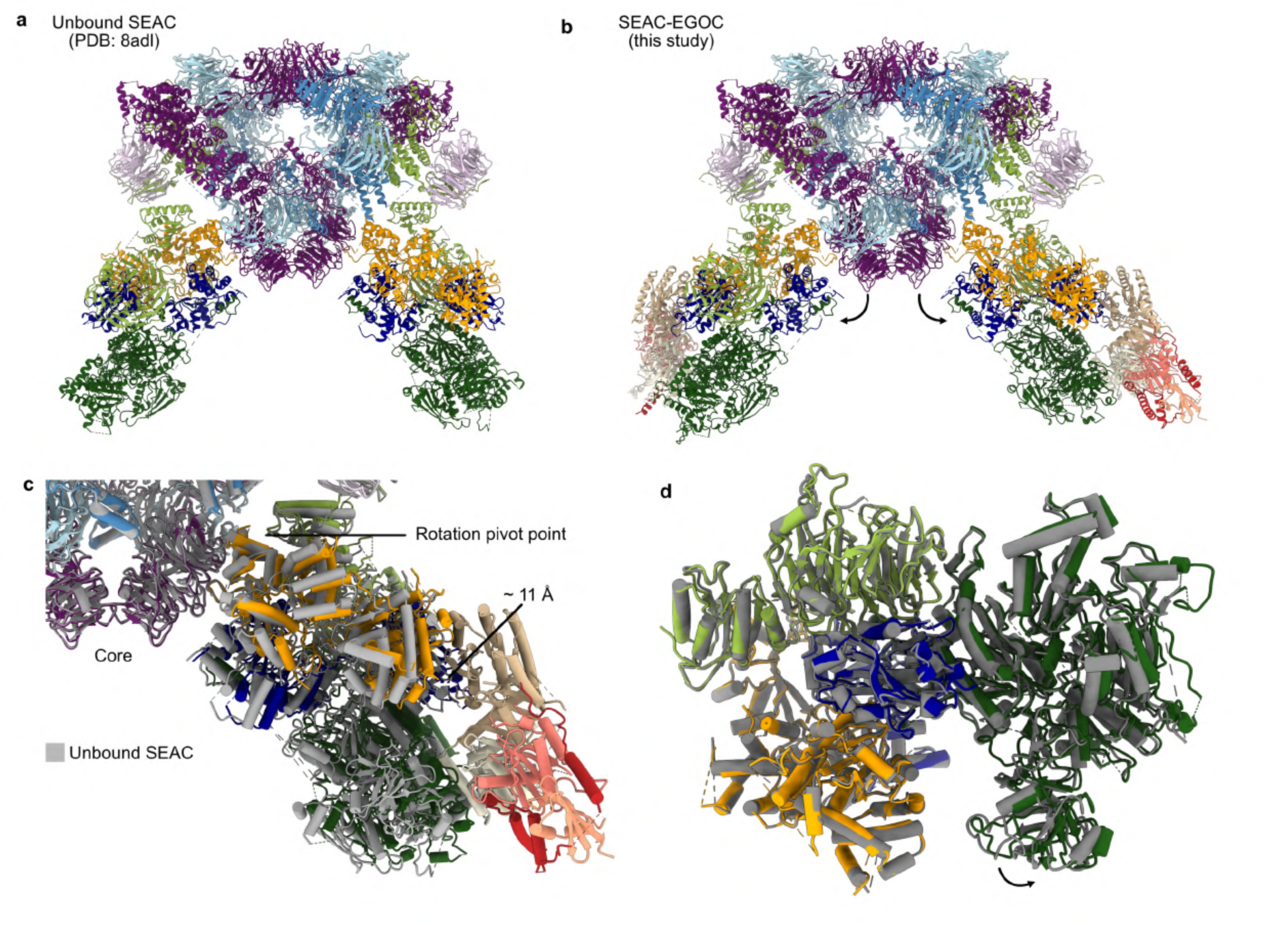
Comparison between unbound and EGOC-bound SEAC structures. **a**, Structure of the unbound SEAC (PDB: 8adl)^10^. **b**, Structure of the EGOC-bound SEAC. Arrows indicate a small opening of the wings relative to the core compared to the unbound structure. **c**, Comparison of the relative position of the SEAC^wing^ between unbound (grey) and EGOC-bound SEAC. The wing rotates around a pivot on Npr3-Sea4, which causes a displacement of the active site of ∼11 Å (distance between Npr2 arginine finger in both models). Models were superimposed on the Sea4 copies that form the back interface, which is adjacent to the wings and interacts with Npr3. **d**, Comparison of the superimposed SEAC^wing^ between unbound and EGOC-bound SEAC. There is a small shift in Sea1 that serves to accommodate Gtr2.

**Extended Data Figure 4.**
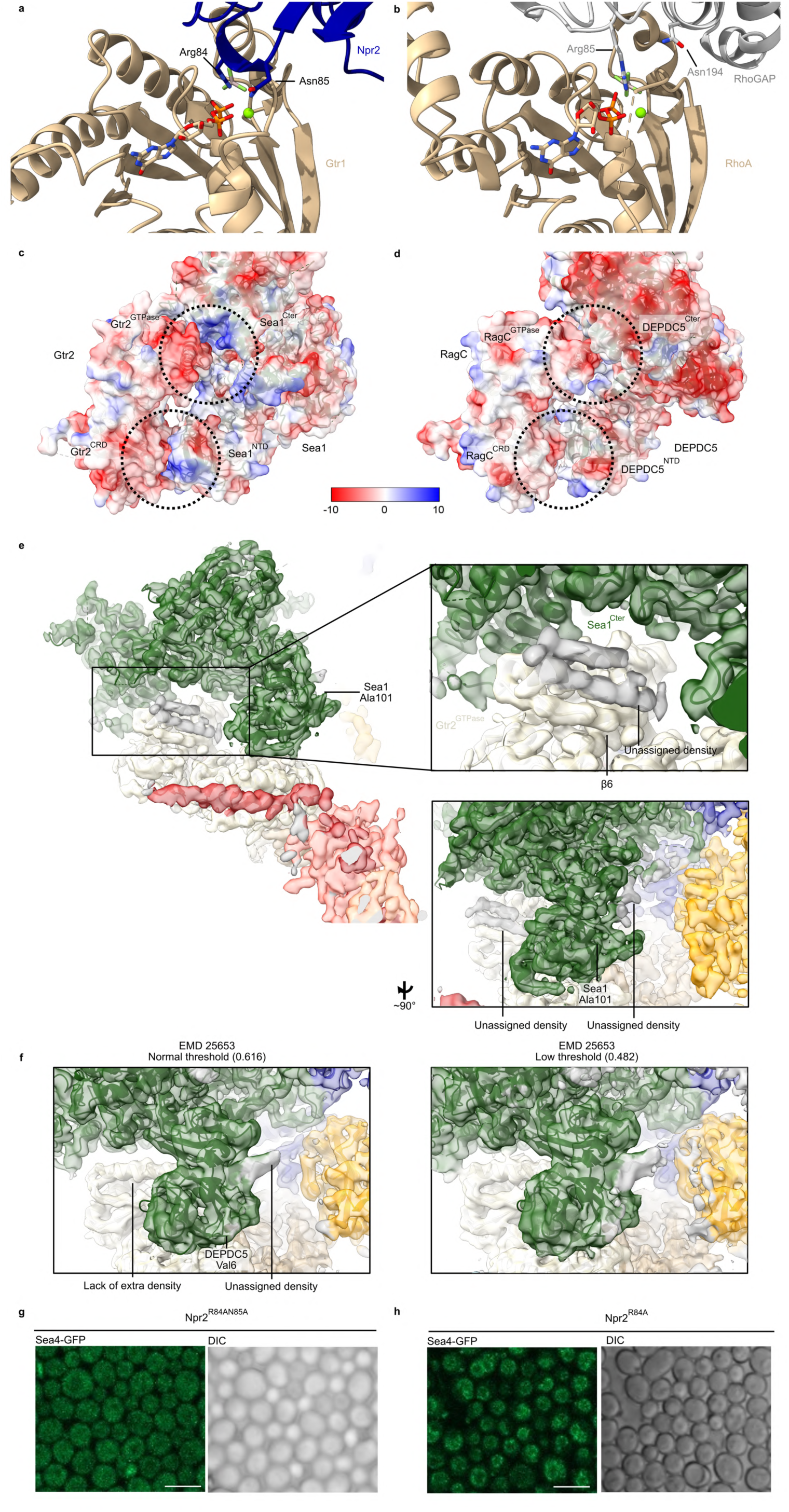
Comparison between the SEAC^wing^, RhoGAP and GATOR1. **a**, Interaction between Npr2 and Gtr1, showing the positions of the arginine finger (Arg84) and neighbouring asparagine (Asn85). **b**, Structure of RhoGAP and RhoA (PDB: 1tx4)^56^, indicating the arginine finger (Arg85) and auxiliary asparagine (Asn194). **c**, Electrostatic potential of the Gtr2-Sea1 binding interface. **d**, Electrostatic potential of the RagC-DEPDC5 binding interface (PDB: 7t3b)^18^. **e**, Unassigned cryo-EM densities between Gtr2 and Sea1 (top panel) and the last visible N-terminal residue of Sea1 (bottom panel, Ala101). Due to the position, the density next to Gtr2 likely corresponds to the disordered first 100 amino acids of Sea1. **f**, Cryo-EM map of the GATOR1-Ragulator-Rag structure (EMD:25653)^18^ showing the absence of density next to Gtr2, even at a lower threshold. This is consistent with the lack of the N-terminal extension in DEDPC5. **g**, Confocal images of Npr2^R84AN85A^ cells expressing *SEA4-GFP*. **h**, Confocal images of Npr2^R84A^ cells expressing *SEA4-GFP*. Scale bar = 5 μm.

**Extended Data Figure 5.**
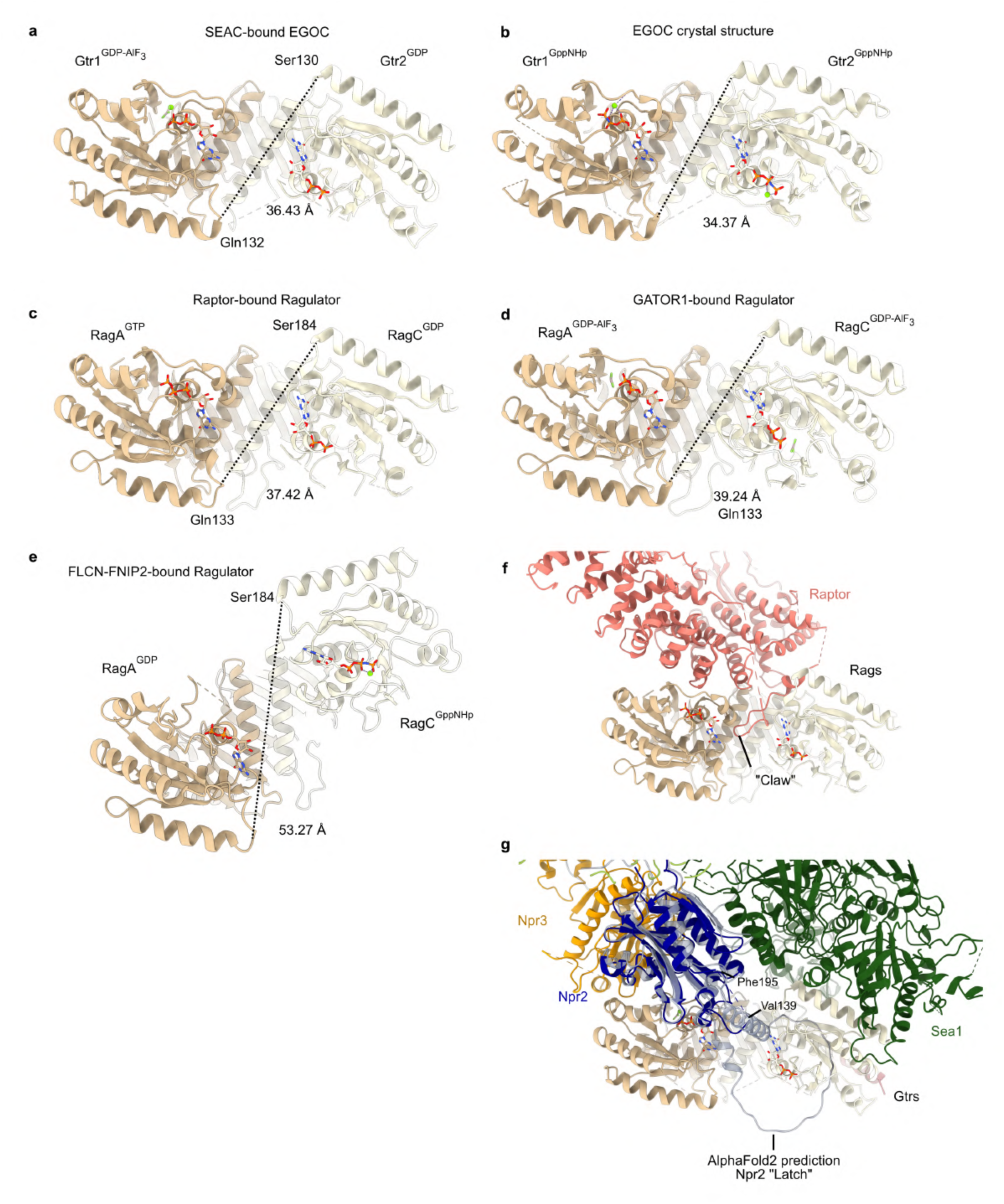
Comparison between Gtr1-Gtr2 and RagA-RagC in different states, and binding of the Rag/Gtr heterodimer to Raptor and SEAC. **a**, Structure of the Gtr1^GDP-AlF3^-Gtr2^GDP^ heterodimer bound to the SEAC. **b,** Structure of Gtr1^GppNHp^-Gtr2^GppNHp^ heterodimer in the EGOC crystal structure (PDB: 6jwp)^1^. **c**, Structure of the RagA^GTP^-Rag^GDP^ heterodimer bound to Raptor (PDB:6u62)^34^. **d**, Structure of the RagA^GDP-AlF3^-RagC^GDP-AlF3^ heterodimer bound to GATOR1 (PDB:7t3b)^18^. **e**, Structure of the RagA^GDP^-Rag^GppNHp^ heterodimer bound to FLCN-FNIP2 (PDB: 6ulg)^29^. The distance between equivalent residues on RagA-RagC and Gtr1-Gtr2 is indicated for each structure, showing that the Gtr heterodimer bound to the SEAC has a very similar conformation as the Raptor-bound Rag heterodimer. Likewise, the inactive RagA-RagC shows a drastically different conformation compared to the other structures. **f**, Structure of Raptor-Ragulator-Rags, with the stretch called the “claw” indicated (PDB: 6u62)^34^. **g**, AlphaFold2 predicted model for Npr2 superimposed onto the SEAC-EGOC structure. Npr2 contains a large, disordered stretch (“Latch”) that is positioned in the vicinity of the arginine finger.

**Extended Data Figure 6.**
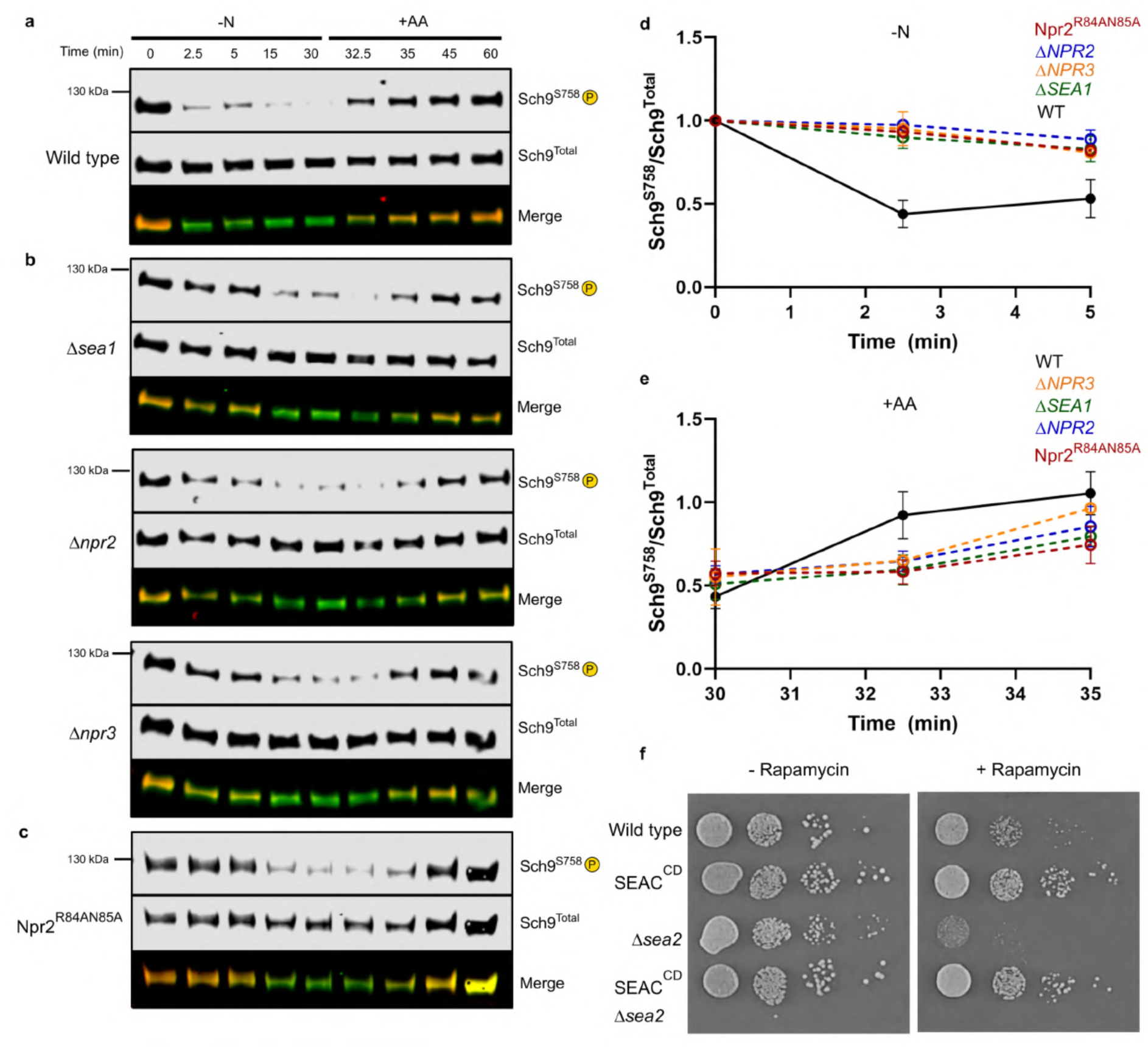
TORC1 amino acid signalling in SEACIT deletion strains. **a**, Immunoblot of phosphorylated and total Sch9 after starvation and repletion of amino acids in wild type cells. **b**, Immunoblot of phosphorylated and total Sch9 after starvation and repletion of amino acids in *Δsea1*, *Δnpr2* and *Δnpr3* cells. **c**, Immunoblot of phosphorylated and total Sch9 after starvation and repletion of amino acids in Npr2^R84AN85A^ cells. **d**, Quantification of relative Sch9 phosphorylation in the first five minutes of starvation in wild type and SEACIT deletion strains. Data from three independent experiments, except for *Δnpr3* (two independent experiments). **e**, Quantification of relative Sch9 phosphorylation in the first five minutes of repletion in wild type and SEACIT deletion strains. Data from three independent experiments, except for *Δnpr3* (two independent experiments). **f**, Growth assays with and without rapamycin (2 nM) for wild type, SEAC^CD^, *Δsea2* and SEAC^CD^*Δsea2* cells

**Extended Data Table 1.**
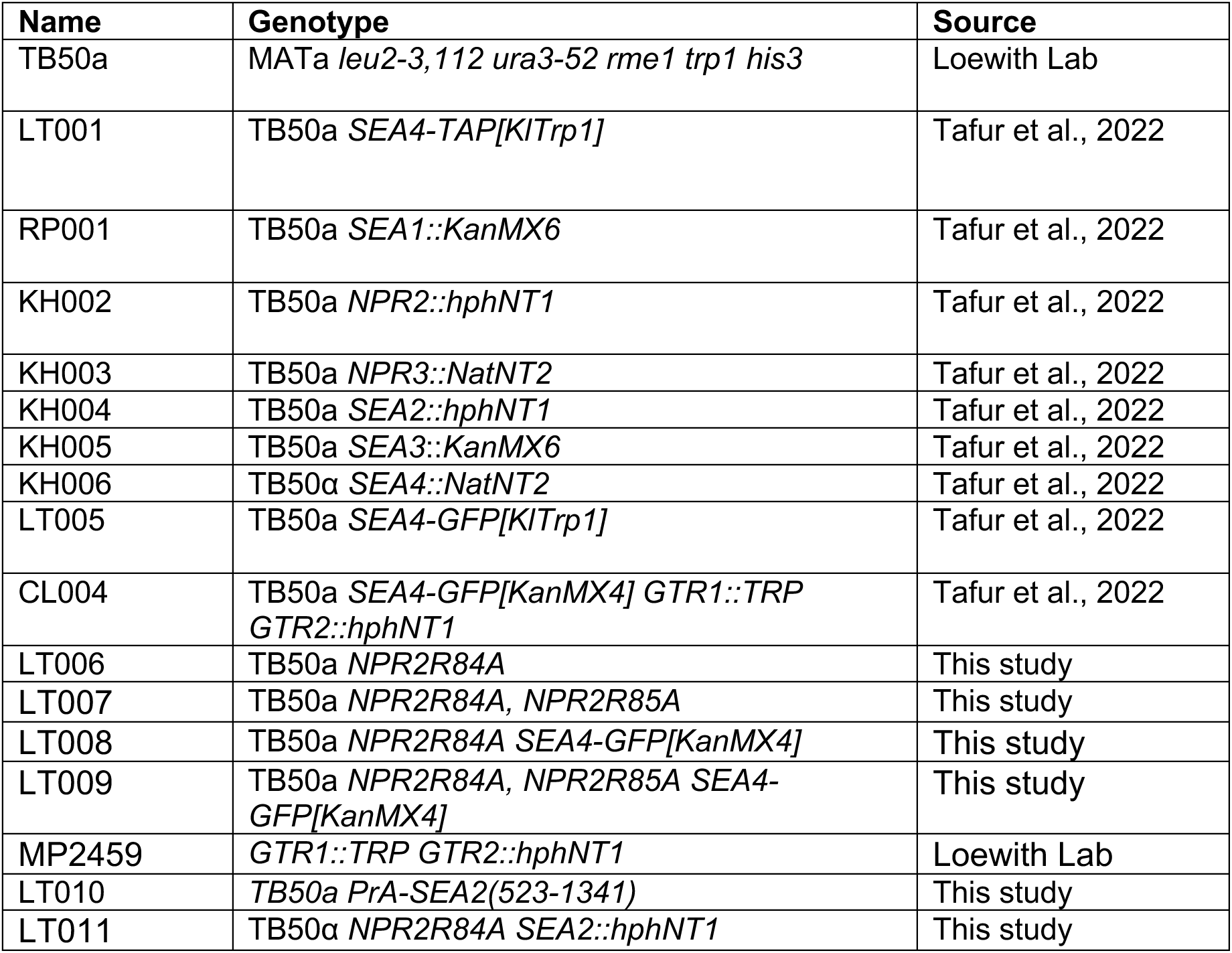
Yeast strains used in this study.

**Extended Data Table 2.** Plasmids used in this study.

| **Plasmid** | **Description** | **Source** |
| --- | --- | --- |
| pST51-Ego2-Ego3-HIS-Ego1 | *pST51; EGO2, EGO3, 6xHIS-EGO1* | This study |
| pST93-Gtr1-Gtr2 | *pST93; GTR1, GTR2* | Tafur et al., 2022 |
| pMB1483 | *pRS415; GTR1-Q65L* | Binda et al., 2009 |
| pJU656 | *pRS415; GTR1-S20L* | Binda et al., 2009 |
| pJU655 | *pRS416; GTR2-Q66L* | Binda et al., 2009 |
| pJU654 | *pRS416; GTR2-S23L* | Binda et al., 2009 |

